# Diagnostic histone modification analysis of individual preimplantation embryos

**DOI:** 10.1101/2023.07.20.549969

**Authors:** Yiren Zeng, Yoichiro Hoshino, Kazuki Susami, Shinnosuke Honda, Naojiro Minami, Shuntaro Ikeda

## Abstract

**Background:** We previously reported a modification of the CUT&Tag method (NTU-CAT) that allows genome-wide histone modification analysis in individual preimplantation embryos. In the present study, NTU-CAT was further simplified by taking advantage of the Well-of-the-Well (WOW) system, which enables the processing of multiple embryos in a shorter time with less reagent and cell loss during the procedure (WOW-CUT&Tag, WOW-CAT).

**Results:** WOW-CAT allowed histone modification profiling from not only a single blastocyst but also from a portion of it. WOW-CAT generated similar H3K4me3 profiles as NTU-CAT, but they were closer to the profiles produced by chromatin immunoprecipitation-sequencing, such as a valley-like trend, indicating that WOW-CAT may attenuate the bias of Tn5 transposase to cut open chromatin regions. Simultaneous WOW-CAT of two halves of single blastocysts was conducted to analyze two different histone modifications (H3K4me3 and H3K27ac) within the same embryo. Furthermore, trophectoderm cells were biopsied and subjected to WOW-CAT in anticipation of preimplantation diagnosis of histone modifications. WOW-CAT allowed the monitoring of epigenetic modifications in the main body of the embryo. For example, analysis of H3K4me3 modifications of *XIST* and *DDX3Y* in trophectoderm biopsies could be used to sex embryos in combination with quantitative PCR, but without the need for deep sequencing.

**Conclusions:** These results suggest the applicability of WOW-CAT for flexible epigenetic analysis of individual embryos in preimplantation epigenetic diagnosis.

## Background

Genome research has been revolutionized by the availability of massively parallel sequencing and its dramatically reduced costs, so that the current difficulties are mostly related to the development of methods to obtain high-quality chromatin fragments for sequencing (1). Epigenomic profiling of preimplantation embryos may be useful for locating abnormal epigenetic modifications and revealing the reason for compromised embryo quality caused by adverse environmental effects that vary according to the conditions and protocols for embryo production such as ovarian stimulation and *in vitro* production (2–6). Temporally and spatially appropriate epigenetic modifications play a crucial role in gene expression in preimplantation embryos (7).

As the last stage before embryo transfer, the quality of blastocysts produced *in vitro* requires close attention, and the methods for detecting epigenetic modifications of individual embryos are continually being improved. Genome-wide investigations of post-translational histone modifications in early embryos used to be performed using mainly chromatin immunoprecipitation-sequencing (ChIP-seq) (8–10). However, because ChIP-seq requires a large number of cells, it is difficult to use this approach to analyze a single embryo that usually contains ∼100 cells. To overcome the disadvantages of ChIP-seq to analyze single or small numbers of preimplantation embryos, several methods have been developed such as enzyme-tethering methods for unfixed cells, in which a specific protein of interest is targeted *in situ* (within the cells) and then profiled genome-wide. For example, Cleavage Under Targets and Release Using Nuclease (CUT&RUN), which is based on Laemmli’s Chromatin ImmunoCleavage strategy (11), targets a chromatin protein through successive binding of a specific antibody and a protein A (pA)-micrococcal nuclease fusion protein that cuts and releases the nearby DNA fragments into the reaction supernatant for subsequent adaptor tagmentation (12, 13). Then, Cleavage Under Targets and Tagmentation (CUT&Tag) was developed, in which a fusion protein of Tn5 transposase and pA (pA-Tn5) is used instead of pA-micrococcal nuclease, enabling the cutting and adaptor tagmentation of the fragments *in situ* simultaneously (14, 15).

Nowadays, CUT&Tag is used to profile representative histone modifications in single blastocysts without binding to a solid phase (NON-TiE-UP CUT&Tag [NTU-CAT]), which has made this method more readily applicable (16). After the successful implementation of NTU-CAT, we hypothesized that this method could be simplified and optimized by using the Well-of-the-Well (WOW) system (17, 18), which was named WOW-CUT&Tag (WOW-CAT). With this method, a maximum of 13 embryos could be treated simultaneously in one WOW dish until the tagmentation step by only liquid exchange without touching the embryos, which saved a considerable amount of time and reaction reagents compared with NTU-CAT. The reduction of systematic errors and cell loss with this approach was considered to facilitate the recovery of DNA that could be sequenced and to enable the profiling of histone modifications using not only a single blastocyst but also a small part of it.

## Methods

### *In vitro* production of bovine blastocysts

This study was approved by the Animal Research Committee of Kyoto University (permit numbers R3-10, R4-10, and R5-10) and was conducted in accordance with the Regulations on Animal Experimentation at Kyoto University. The bovine ovaries used in this study were purchased from a commercial abattoir as by-products of meat processing, and the frozen bull semen (conventional and Y chromosome-sorted) used for *in vitro* fertilization (IVF) was also commercially available. *In vitro* production of bovine embryos by IVF was performed as previously described (19, 20). Blastocyst-stage embryos at 168–192 h post-IVF (days 7–8) were collected individually.

### Blastocyst biopsy

Blastocyst biopsy was performed as described by de Sousa et al. (21) with minor modifications. Briefly, blastocysts were cut using a micromanipulator (Leica Microsystems, Wetzlar, Germany) and a stainless-steel blade at an angle of 30° (Bio-Cut-Blades Feather; Feather Safety Razor Co., Osaka, Japan). Embryos were micromanipulated on a 90 × 15 mm Petri dish (AS ONE Corporation, Osaka, Japan) containing 200 µL holding medium consisting of TCM–199 and Hanks’ salts (Invitrogen, Carlsbad, CA) supplemented with 0.3% bovine serum albumin (BSA; Sigma-Aldrich, St. Louis, MO).

### WOW-CAT

For whole blastocysts, the zona pellucida was freed from the embryos by 0.5% (w/v) pronase treatment and washed with phosphate-buffered saline (PBS) containing 0.01% (w/v) polyvinyl alcohol (PVA) and 1% (v/v) Protease Inhibitor Cocktail (PIC) (PBS-PVA-PIC) before allocation into the WOW dish (LinKID micro25 Culture Dish; Dai Nippon Printing, Tokyo, Japan). The biopsies and remaining parts of the blastocysts were washed with PBS-PVA-PIC before placing in the WOW dish. Then, individual samples were transferred into separate microwells in the WOW dish by individual pipettes to avoid cross-contamination and all the treatments onward were conducted in one WOW dish until pA-Tn5 binding by only liquid exchange. The basal kit for CUT&Tag was a CUT&Tag-IT Assay Kit (Active Motif, Carlsbad, CA). Dig-Wash buffer and Dig-300 buffer in the kit were supplemented with 0.01% (w/v) PVA to avoid cell adhesion to the wall of the dishes or pipettes. Primary antibody binding was performed in the WOW dish with 100 µL Antibody Buffer, which contained 2 µL (2.8 and 5.6 µg for H3K4me3 and H3K27ac, respectively) of the primary antibodies (C15410003 and C15410196 for H3K4me3 and H3K27ac, respectively; Diagenode, Denville, NJ), 0.05% (w/v) digitonin, and 1% (v/v) PIC. The blastocysts were incubated at 4°C overnight with gentle shaking (400 rpm). A negative control was set by omitting the primary antibodies. After the primary antibody reaction, the primary antibody buffer was replaced with 100 µL Dig-Wash buffer containing a secondary antibody (guinea pig anti-rabbit IgG antibody, 1 μL), 0.05% (w/v) digitonin, and 1% (v/v) PIC and incubated for 1 h at room temperature (400 rpm). After three washes with 100 µL Dig-Wash Buffer supplemented with digitonin and PIC, pA-Tn5 binding was performed using 100 µL Dig-300 Buffer replacement, which contained 1 μL pA-Tn5 transposomes, 0.01% (w/v) digitonin, and 1% (v/v) PIC and incubated for 1 h at room temperature (400 rpm). After three washes with 100 µL Dig-300 Buffer supplemented with digitonin and PIC, the cells were transferred to individual microcentrifuge tubes (Eppendorf 0030 108.051) containing 125 µL Tagmentation Buffer with 0.01% (w/v) digitonin and 1% (v/v) PIC and incubated for 1 h at 37 °C without shaking.

After tagmentation, 4.2 μL of 0.5 M EDTA, 1.25 μL of 10% SDS, and 1.1 μL of 10 mg/mL proteinase K were added to each tube and incubated for 1 h at 55 °C with vigorous shaking (1,300 rpm). After cooling to room temperature, SPRIselect beads (145 µL; Beckman Coulter, Brea, CA) were added to each tube, vortexed for 1 min, and allowed to incubate for 10 min at room temperature. The tubes were placed on a magnetic stand for 4 min to collect the magnetic beads and the liquid was removed. The beads were washed twice with 1 mL of 80% ethanol. After drying the bead pellets for 2–5 min, 35 µL DNA Purification Elution Buffer was added and the tubes were vortexed and left to stand for 5 min at room temperature. The tubes were placed on a magnetic stand for 4 min to collect the magnetic beads and the liquid containing tagmented DNA was transferred to PCR tubes.

PCR amplification of sequencing libraries was performed in a volume of 50 µL using 30 µL tagmented DNA and i7 and i5 indexing primers according to the manufacturer’s protocol. The PCR conditions were as follows: 72°C for 5 min; 98°C for 30 s; 20 cycles for blastocysts (23 cycles for biopsies and 21 cycles for the remaining parts or halved blastocysts) of 98°C for 10 s and 63°C for 10 s; final extension at 72 °C for 1 min; and hold at 10°C. Post-PCR library purification was performed with 55 µL SPRIselect beads (vortex for 1 min, stand for 5 min, and bead collection for 4 min) and 180 µL of 80% ethanol as described above. Finally, the sequencing libraries were eluted in 25 µL DNA Purification Elution Buffer.

### Immunofluorescence

To confirm the number and type of cells in the biopsies, the biopsied and remaining parts of the blastocysts were subjected to CDX2 immunolabeling according to the method of Wydooghe et al. (22) with some modifications. Briefly, the biopsied and remaining parts were fixed in 10% (v/v) formalin neutral buffer solution (Fujifilm Wako Pure Chemical Corp., Osaka, Japan) for 1 h, washed in PBS containing 0.05% (v/v) Tween 20 (PBST) for 1 h and subsequently permeabilized with 0.5% (v/v) Triton X-100 in PBS for 1 h at room temperature. After washing in PBST for 1 h at room temperature, the samples were treated with blocking solution (PBST supplemented with 1% [w/v] BSA) for 1 h at room temperature and subsequently incubated in a ready-to-use primary anti-CDX2 antibody solution (BioGenex, Fremont, CA) overnight at 4°C. After washing with blocking solution for 1 h at room temperature, the samples were incubated for 3 h at room temperature in the presence of Alexa Fluor 546-conjugated goat anti-mouse IgG (1:1,000; Life Technologies, Carlsbad, CA). Nuclei were counterstained with 10 μg/mL Hoechst 33342 in PBST for 20 min. The samples were washed with PBST, mounted on slides with a droplet of VECTASHIELD mounting medium (Vector Laboratories, Burlingame, CA) and flattened with a coverslip. The slides were examined under a fluorescence microscope. The total number of cells was counted based on the Hoechst image, and the number of trophectoderm (TE) cells, which had been stained by both the anti-CDX2 antibody and Hoechst, was determined based on the merged images. The number of inner cell mass (ICM) cells was calculated by subtracting the number of TE cells from the total number of cells.

### WOW-CAT-qPCR

Quantitative PCR (qPCR) was conducted to assess the enrichment of specific histone modifications in the WOW-CAT libraries. The qPCR mixture (total volume: 10 µL) was prepared as follows: 5 µL THUNDERBIRD Next SYBR qPCR (Toyobo, Osaka, Japan), 1 µL library DNA, and 0.3 µL (10 µM) forward/reverse primers. PCR was performed with a StepOnePlus Real-time PCR system (Life Technologies) using the following program: 95°C for 30 s, followed by 40 cycles at 95°C for 5 s and 60°C for 10 s with melt curve drawing. The primers used are listed in Table 1.

**Table 1.**
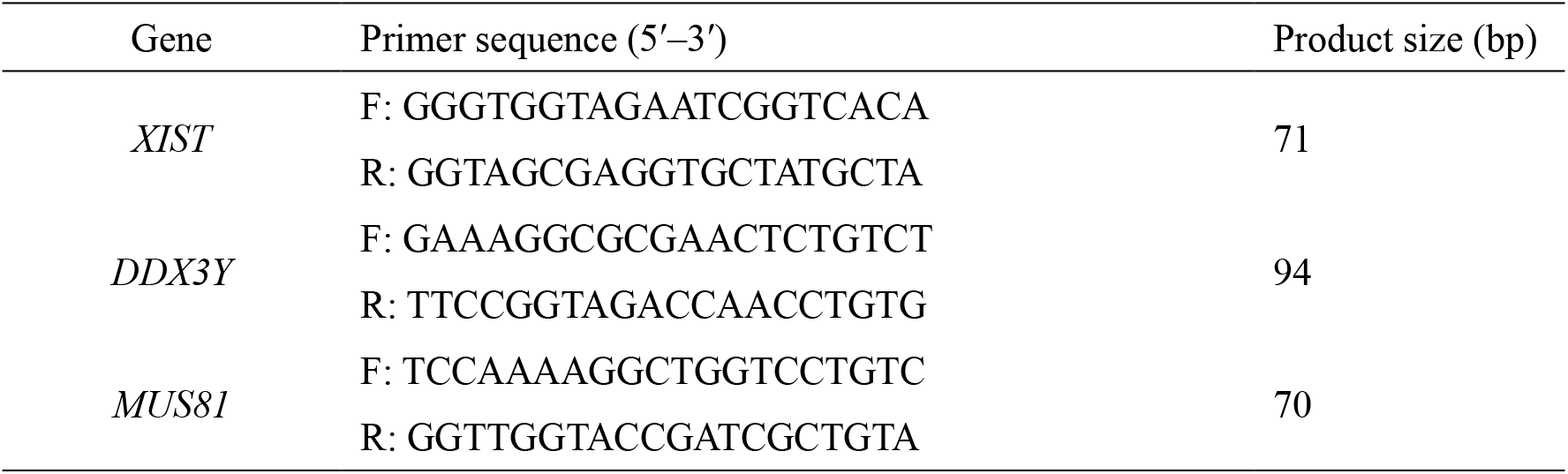
Primers used for WOW-CAT-qPCR for sex identification.

### Sex identification

We examined specific histone modifications that could be used for sex determination of the embryos. We focused on H3K4me3 modifications of the *XIST* and *DDX3Y* genes as putative female- and male-specific markers, respectively, from the WOW-CAT-next-generation sequencing (NGS) results. We conducted two experiments as follows. First, we biopsied the blastocysts as described above, and the biopsies were subjected to WOW-CAT-qPCR for *XIST* and *DDX3Y*. *MUS81* was used as an internal control. The remaining parts paired with the biopsies were subjected to conventional PCR-based sex identification using Y chromosome-specific repeat sequences as previously described (23, 24). Then, we assessed the consistency of both methods. Second, to further examine the reliability of H3K4me3 modifications at *XIST* and *DDX3Y* as female- and male-specific markers, respectively, IVF with Y chromosome-sorted sperm was conducted, and the derived blastocysts were subjected to WOW-CAT-qPCR to detect H3K4me3 at *XIST* and *DDX3Y*.

### DNA sequencing and data processing

Sequencing and data processing were performed as previously reported (16). Briefly, paired-end 150-base pair sequencing reads generated by a HiSeqX (Illumina) were quality checked, merged, and aligned to the bovine genome (Bos_taurus_UMD_3.1.1/bosTau8, June 2014) using Bowtie 2 (25). Handling of sam and bam files was performed by Samtools (http://www.htslib.org/). Mapping duplicates were removed by Picard (http://broadinstitute.github.io/picard/). The generated bam files were converted to bigWig files by the bamCoverage tool of deepTools (https://deeptools.readthedocs.io/en/develop/) with counts-per-million normalization. The correlation plots between the experiments were made from bigWig files fed to deepTools. Average H3K4me3 signal profiles were generated by ngs.plot (26). Peaks were visualized using Integrative Genomics Viewer (27).

### Publicly available data

For the comparison with the present WOW-CAT results, we used ChIP-seq (rep1 and rep3 of GSE16122) (19) and NTU-CAT (Rep1–4 of https://zenodo.org/record/6002122) (16) data, respectively.

## Results

### Schema for WOW-CAT

The schema for WOW-CAT is shown in Fig. 1. After removal of the zona pellucida or biopsy, the individual embryos or cell masses were transferred into separate microwells in the WOW dish containing a primary antibody solution with a detergent (digitonin) and incubated. Although we had 25 microwells per dish, the maximum number of embryos processed per dish was 13 because the wells were used so that samples were not adjacent to each other vertically and horizontally. Subsequent secondary antibody reactions and the tethering of pA-Tn5 fusion protein with sequencing adapters were performed by exchanging the respective reaction solutions in the WOW dish. The tagmentation reaction with pA-Tn5 activation was performed in microtubes and the tagmented DNA was extracted and PCR amplified using index primers, and the library after purification was used for sequencing.

**Figure 1.**
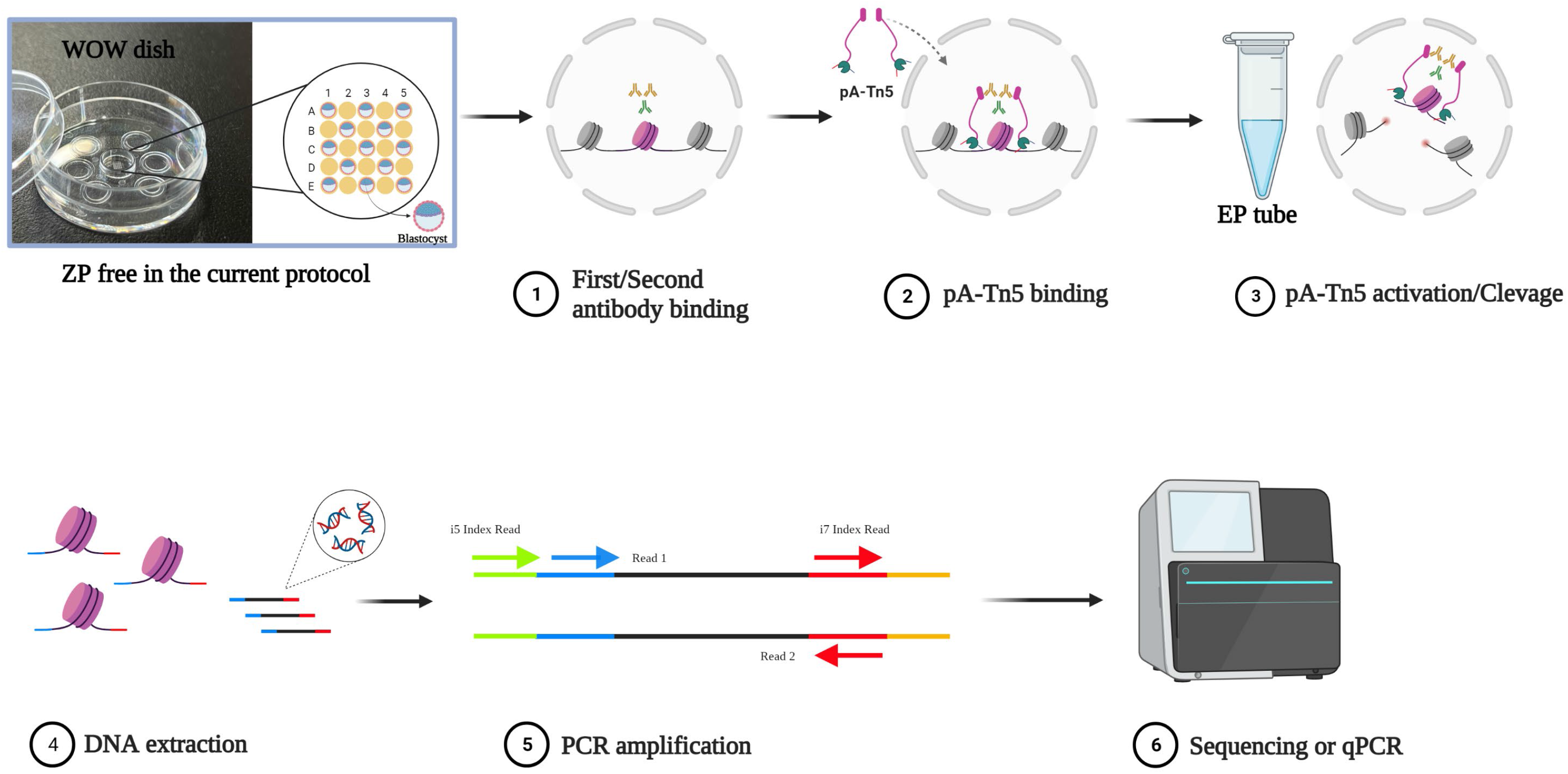
Schematic diagram of WOW-CAT. See the “Methods” section for details. All icons except the WOW dish picture are from Biorender.com.

### H3K4me3 profile of whole blastocysts assessed by WOW-CAT

Since we had previously validated NTU-CAT and obtained comparable results to ChIP-seq for the H3K4me3 modification (16), we initially targeted this modification for WOW-CAT application. A snapshot of the WOW-CAT peaks for the H3K4me3 modification from four replicates (i.e., four single blastocysts) is shown in Fig. 2a, alongside the peaks from our previous ChIP-seq (19) and NTU-CAT (16) analyses of blastocysts. The overall landscape depicted by the location and shape of the peaks was very similar among the three methods, except for subtle differences in the shapes of the peaks (Fig. 2a). Fig. 2b shows the average profile plots of the H3K4me3 signal around the transcription start sites (TSSs) in these experiments. A striking difference between the NTU-CAT and ChIP-seq profiles was that the valley-like shapes near TSSs detected in ChIP-seq were not detected in NTU-CAT, which was in line with the findings of other studies on CUT&Tag (28, 29). In contrast, WOW-CAT captured a little more of the valley-like shapes, although not as much as ChIP-seq. Pairwise comparisons of H3K4me3 signals showed a high correlation among the methods and replicates (Fig. 2c).

**Figure 2.**
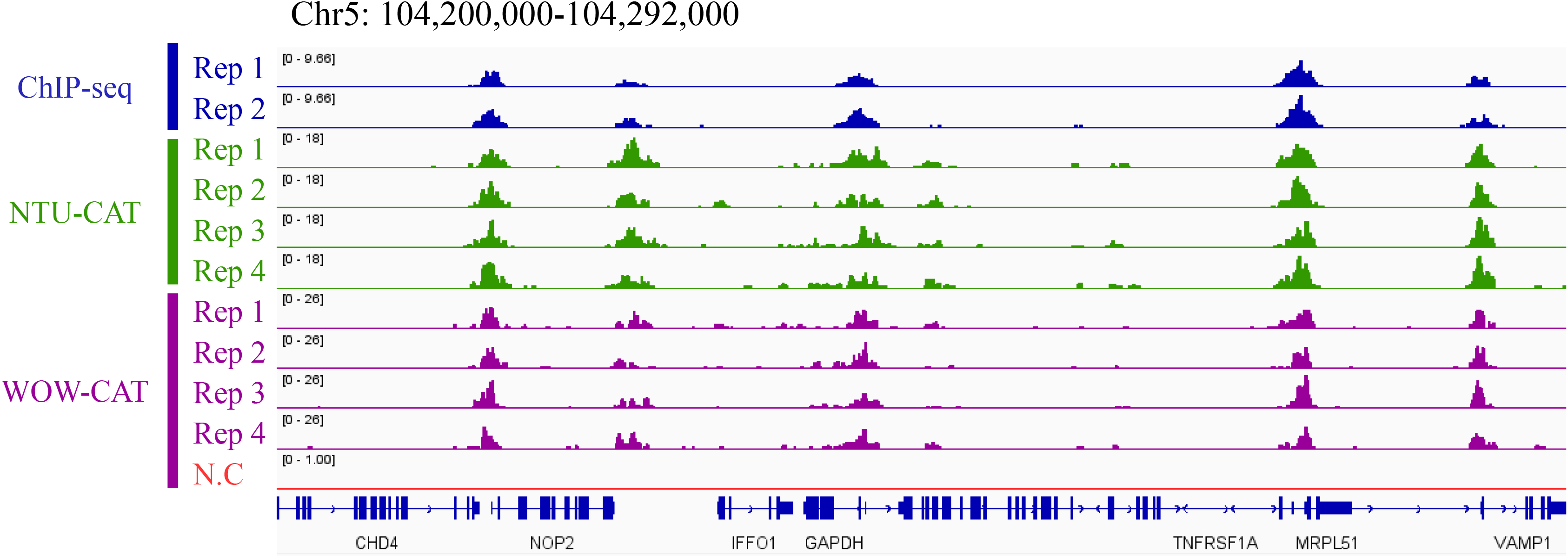
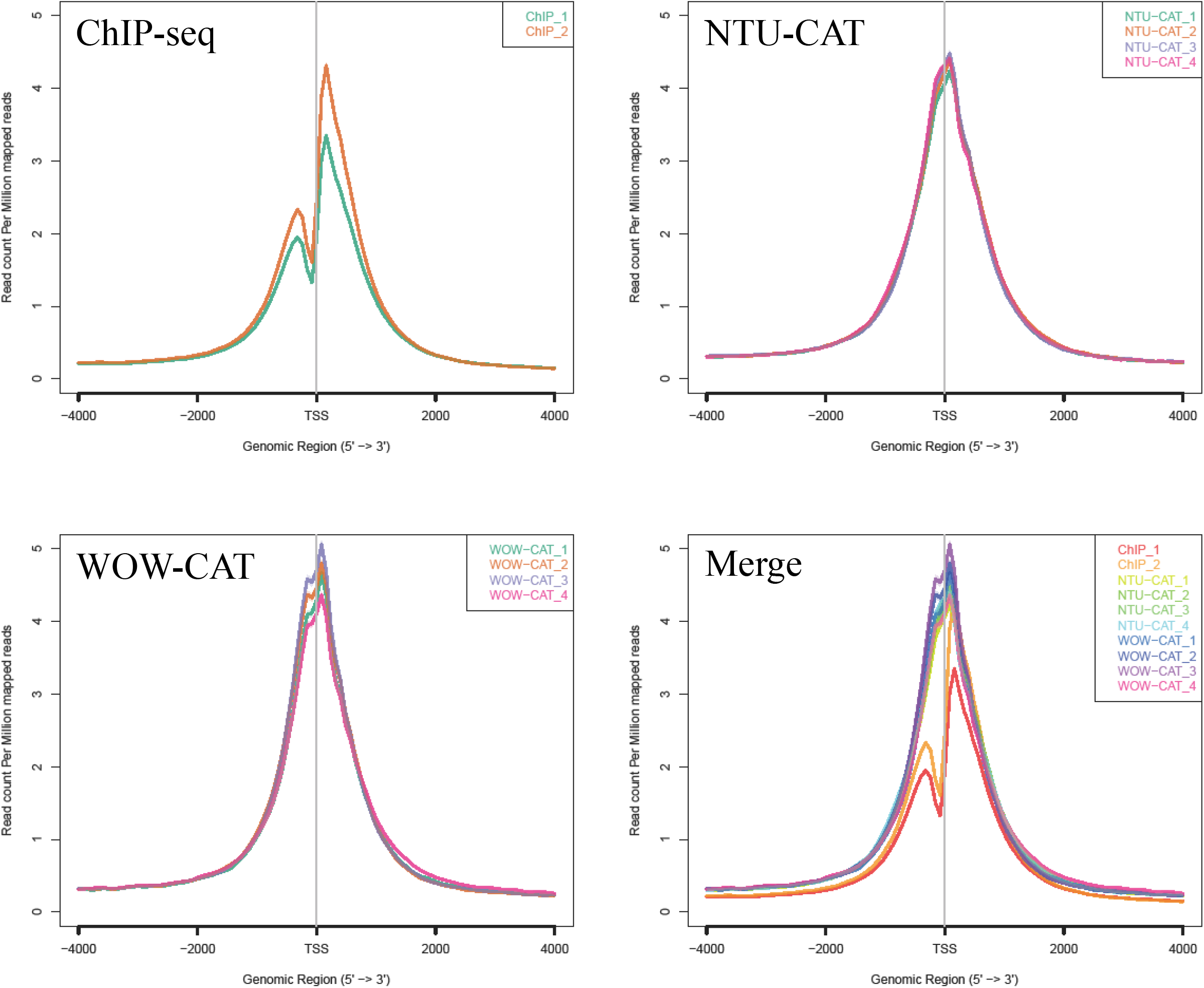
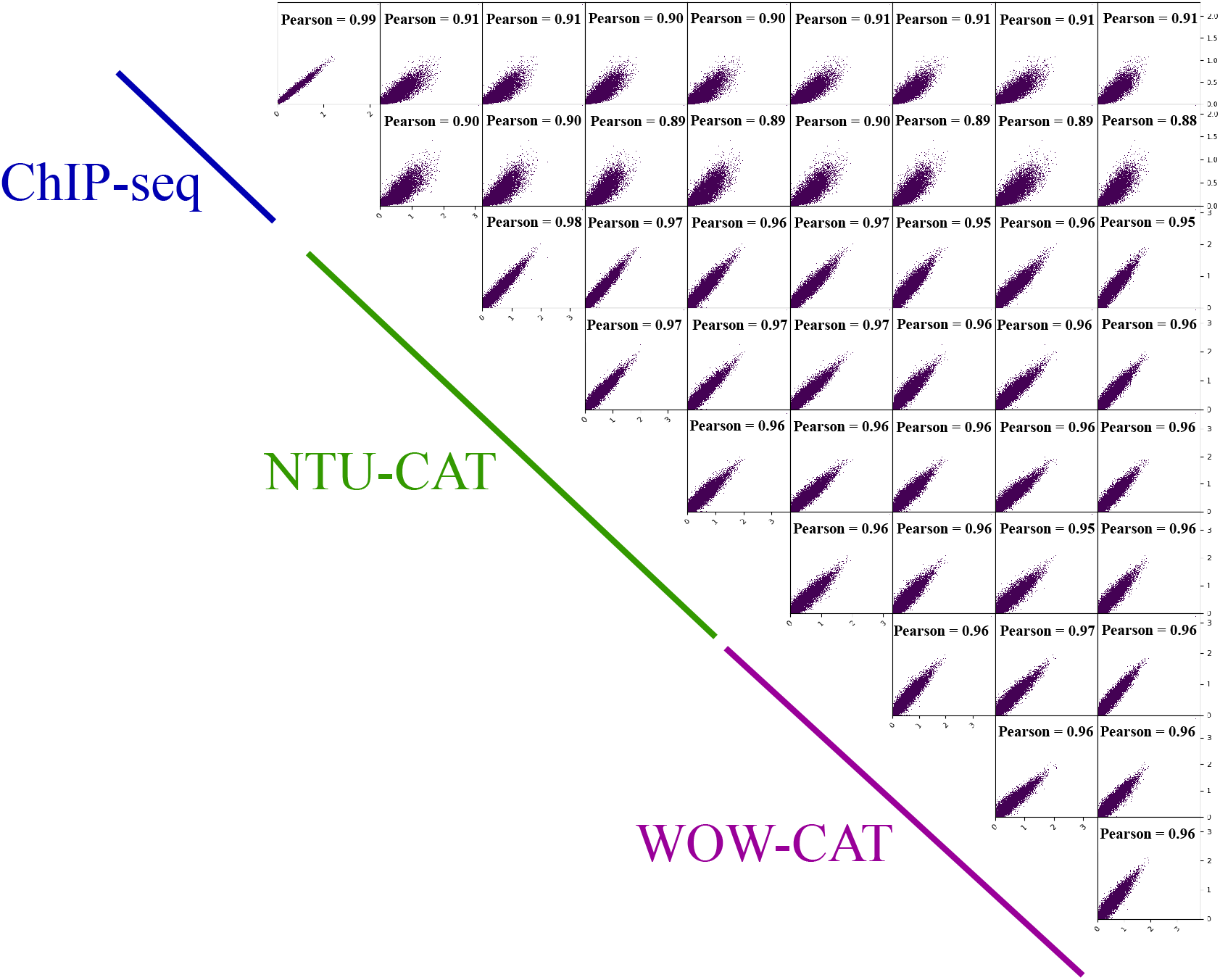
Comparison of WOW-CAT, NTU-CAT, and ChIP-seq results for H3K4me3 in bovine blastocysts. (a) A snapshot of the WOW-CAT peaks from four replicates, alongside the peaks from our previous ChIP-seq and NTU-CAT analyses of blastocysts for the H3K4me3 modification. (b) The average profile plots of H3K4me3 signals around the TSSs in these experiments. (c) Pairwise comparisons of H3K4me3 signals among the different methods and replicates. Scatterplots of these pairwise comparisons are shown with Pearson correlation coefficients. Bin sizes of 10 kb and natural log transformation after adding 1 were used for drawing with deepTools (https://deeptools.readthedocs.io/en/develop/).

### Simultaneous H3K4me3 and H3K27ac profiling within single blastocysts assessed by WOW-CAT

WOW-CAT allows the reaction solution to be changed without moving the cells while they remain at the bottom of the dish, making it easier to complete the immuno-tethering process without losing the samples, even with smaller cell masses. Therefore, we performed the simultaneous profiling of two different histone modifications within the identical single blastocysts, which has never been reported. To accomplish simultaneous profiling, a whole blastocyst was cut equally into two parts, so that each half contained the ICM and TE (Fig. 3a). Then, each half was subjected to WOW-CAT to profile the H3K4me3 and H3K27ac modifications, respectively. Fig. 3a shows the same region as reported in the preprint by Zhou et al. (30), who used 1,000–1,500 blastomeres for CUT&Tag for these modifications. Our present results were similar to those of Zhou et al. in terms of the common (unshaded), H3K4me3-dominant (blue-shaded), and H3K27ac-dominant (red-shaded) areas. The average profile plots of the H3K4me3/H3K27ac signals around the TSSs and gene bodies are shown in Fig. 3b. Highly similar but different modification profiles between H3K4me3 and H3K27ac around the TSSs were detected in this experiment. Fig. 3c and d show the heatmap and volcano plot, respectively, of the normalized H3K4me3 and H3K27ac modifications within three single embryos. These analyses clearly demonstrated H3K4me3- or H3K27ac-dominant genes even within the same embryo (e.g., *NDUFA3* and *GJB3*, respectively) (Fig. 3e). The genes showed enrichment in the biological processes exclusively related to gene expression and signal transduction, respectively (Fig. 3f).

**Figure 3.**
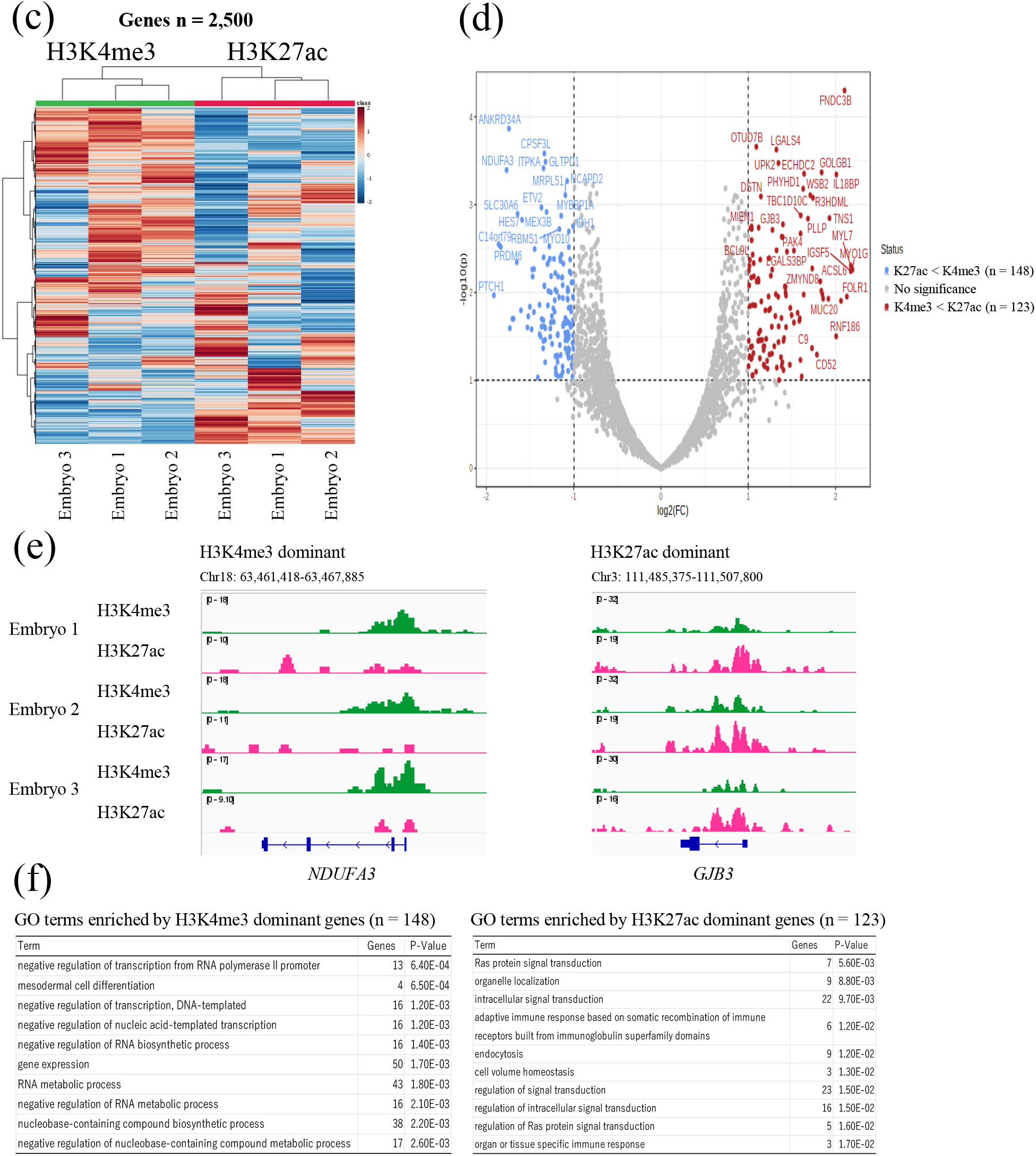
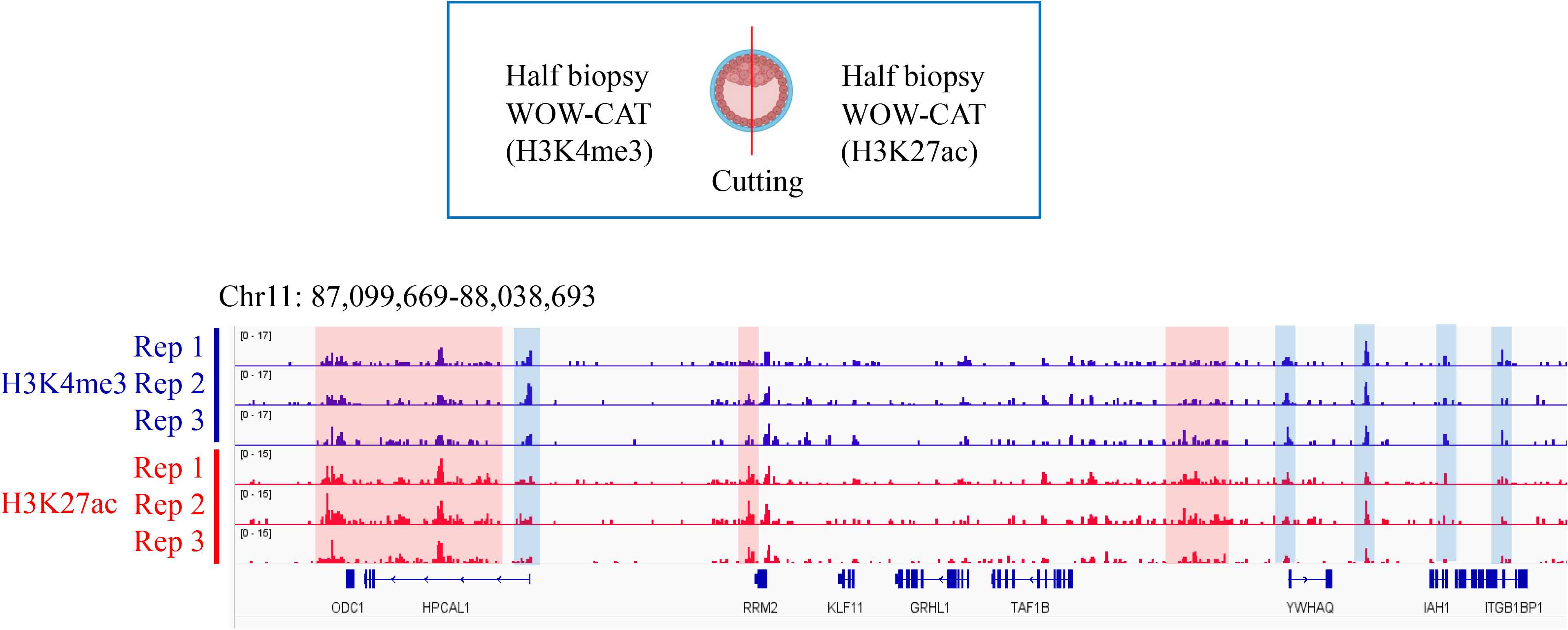
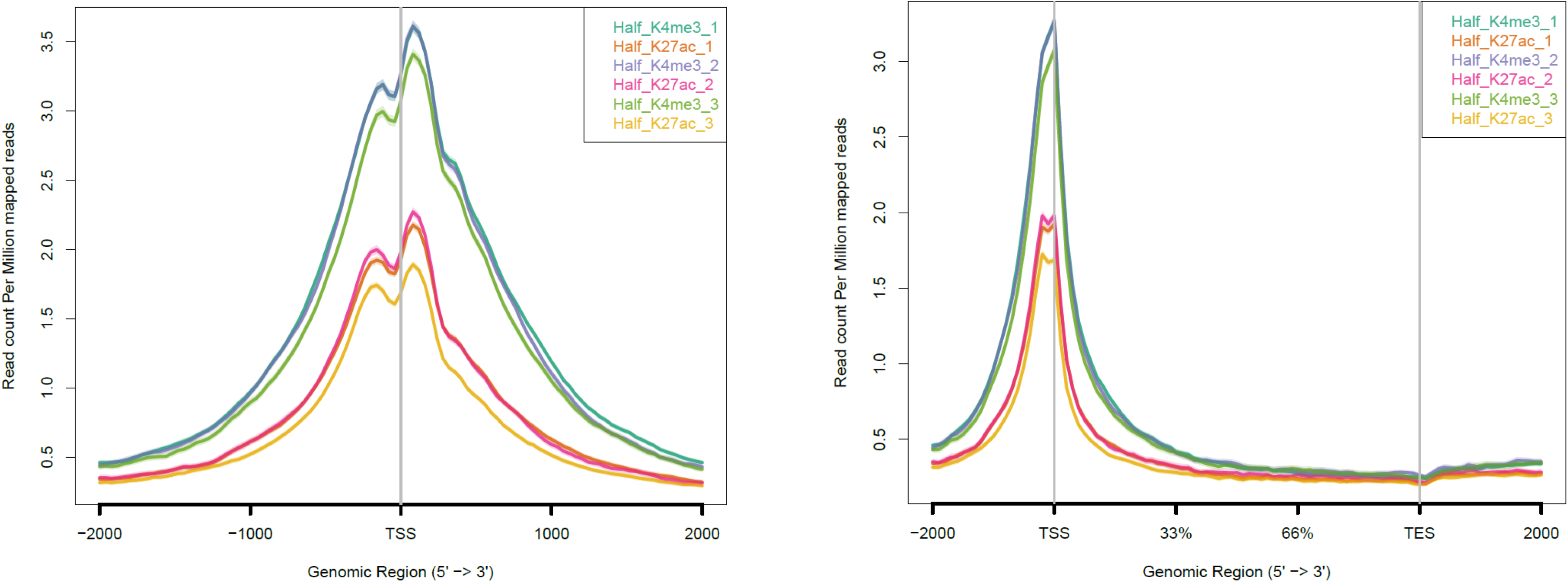
Simultaneous profiling of H3K4me3 and H3K27ac within the same blastocysts assessed by WOW-CAT. (a) The results were similar to those reported by Zhou et al. (30) in terms of the common (unshaded), H3K4me3-dominant (blue-shaded), and H3K27ac-dominant (red-shaded) areas, respectively. Blastocyst icon is from Biorender.com. (b) Average profile plots of H3K4me3/H3K27ac signals around TSSs and gene body regions. (c, d) Heatmap and volcano plot, respectively, of the normalized H3K4me3 and H3K27ac modifications within three single blastocysts. (e) Examples of H3K4me3- and H3K27ac-dominant genes. (f) Gene Ontology terms enriched by H3K4me3- and H3K27ac-dominant genes.

### H3K4me3 profile of biopsied single blastocysts assessed by WOW-CAT

To preserve the integrity of blastocysts for future embryo transfer, we biopsied only as few TE cells as possible to perform WOW-CAT. The precision of the TE cell biopsy was examined by immunofluorescent staining of CDX2 as a TE cell-specific marker in combination with Hoechst 33342 staining of total cell nuclei (Fig. 4a). The high rate of average CDX2/Hoechst-stained cells (21 in 22, 95%) in the biopsied part indicated that the TE cells were precisely excised. In addition, the absence of H3K4me3 modifications of *NANOG* (an ICM-specific marker) in the biopsied part further showed the precision of the TE cell biopsy (Fig. 4b). A snapshot of the WOW-CAT H3K4me3 peaks of the whole, biopsied, and remaining (main) parts of the blastocysts is shown in Fig. 4c. The overall landscape depicted by the location and shape of the peaks was similar among the different samples.

**Figure 4.**
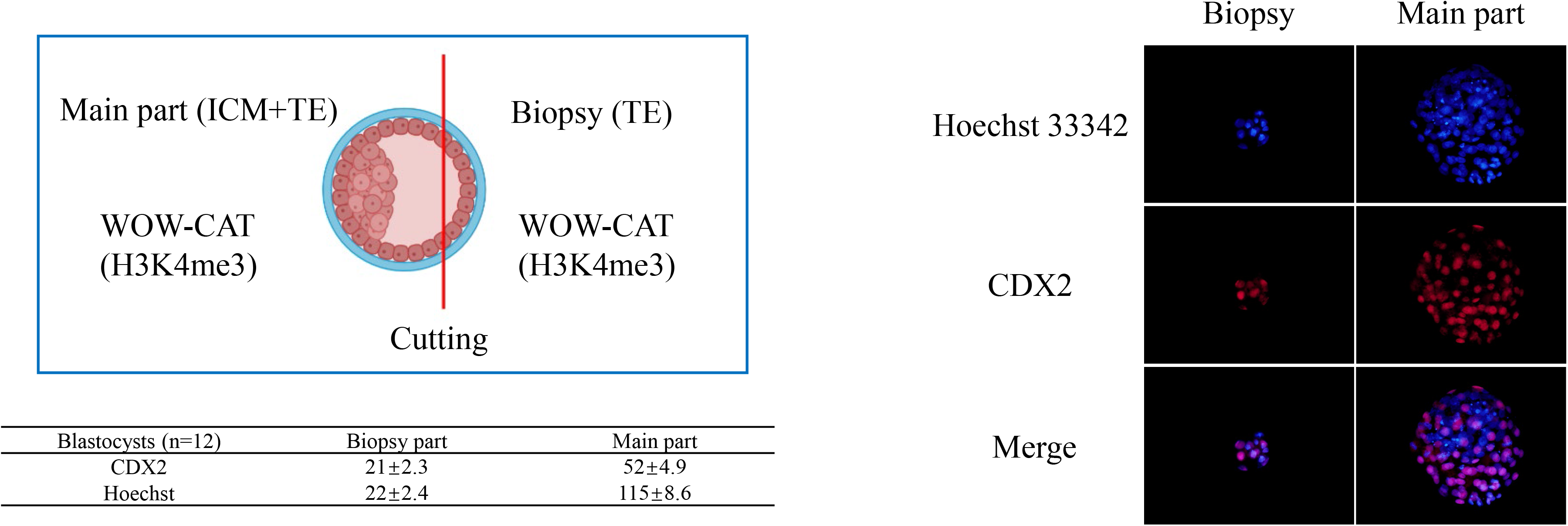
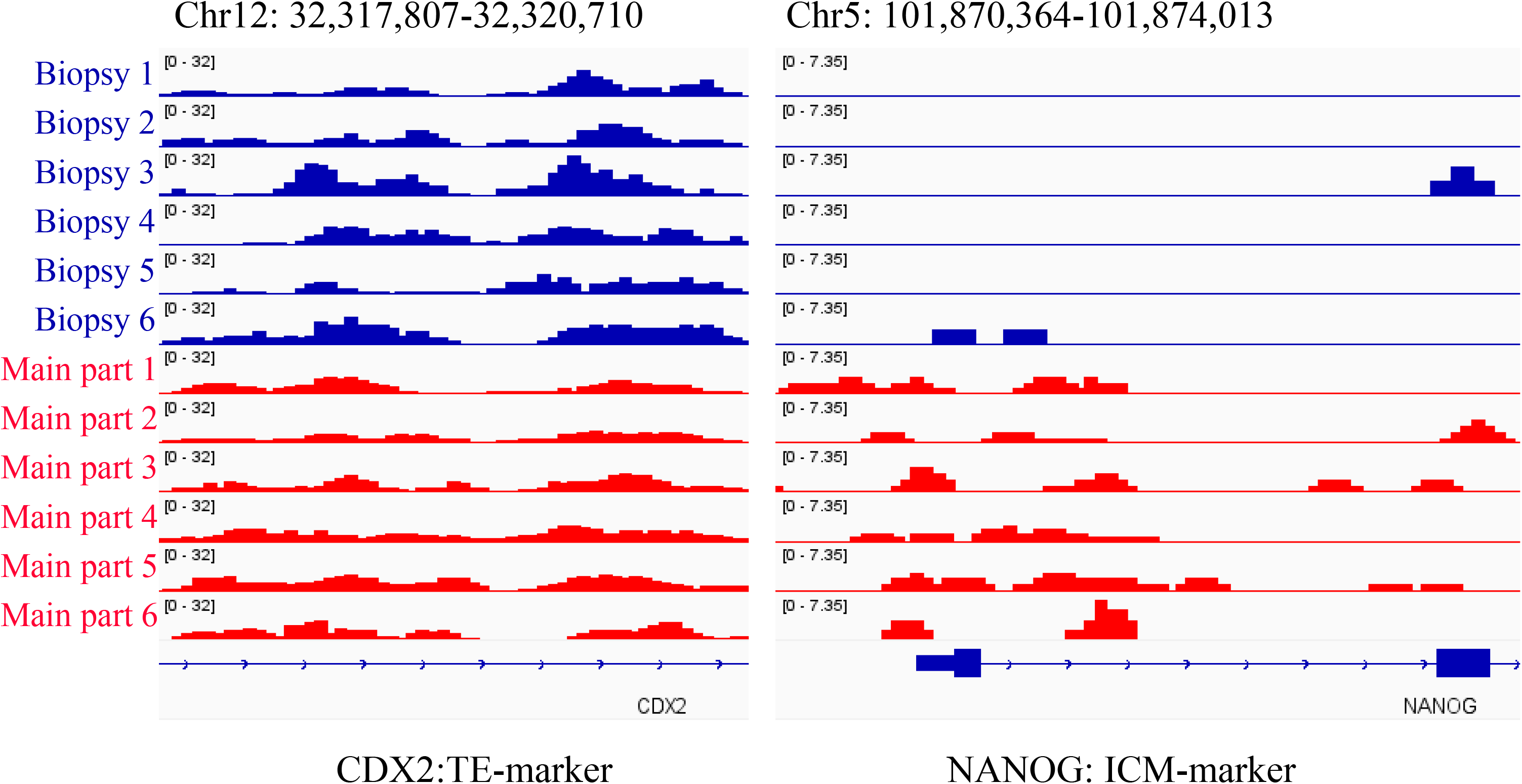
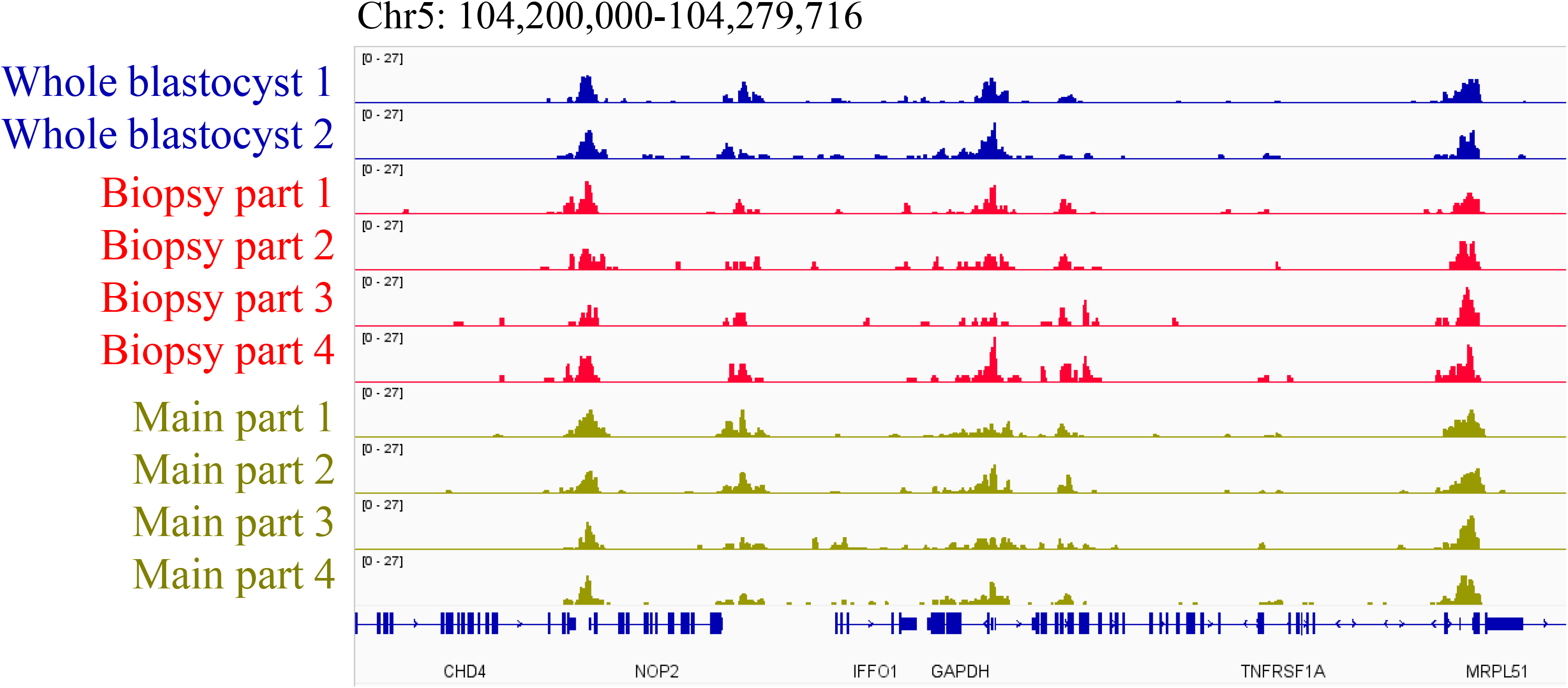
H3K4me3 profile of biopsied single blastocysts assessed by WOW-CAT. (a) The precision of TE cell biopsy examined by immunofluorescent staining of *CDX2* as a TE cell-specific marker in combination with Hoechst 33342 staining of total cell nuclei. The table shows the cell numbers of each part (mean ± standard error of the mean). Blastocyst icon is from Biorender.com. (b) The absence of H3K4me3 modifications at *NANOG* (an ICM-specific marker) in the biopsied parts. (c) A snapshot of the WOW-CAT H3K4me3 peaks of the whole, biopsied, and remaining (main) parts of blastocysts.

### Sex identification: preliminary application of WOW-CAT for embryonic diagnosis

With the successful construction of DNA libraries for the enrichment of specific histone modifications from biopsied TE cells by WOW-CAT, we developed a qPCR-based protocol to detect important histone markers from these libraries, which could directly reflect the properties of the remaining part of the blastocyst. We selected H3K4me3 modifications at *XIST* and *DDX3Y* as male- and female-specific sex identification markers, respectively, from the results of NGS-based WOW- CAT (Fig. 5). Figure 5 shows that most embryos had only one modification, either *XIST* or *DDX3Y*, suggesting they were female and male embryos, respectively. We designed primers for these genes at areas with abundant H3K4me3 modifications.

**Figure 5.**
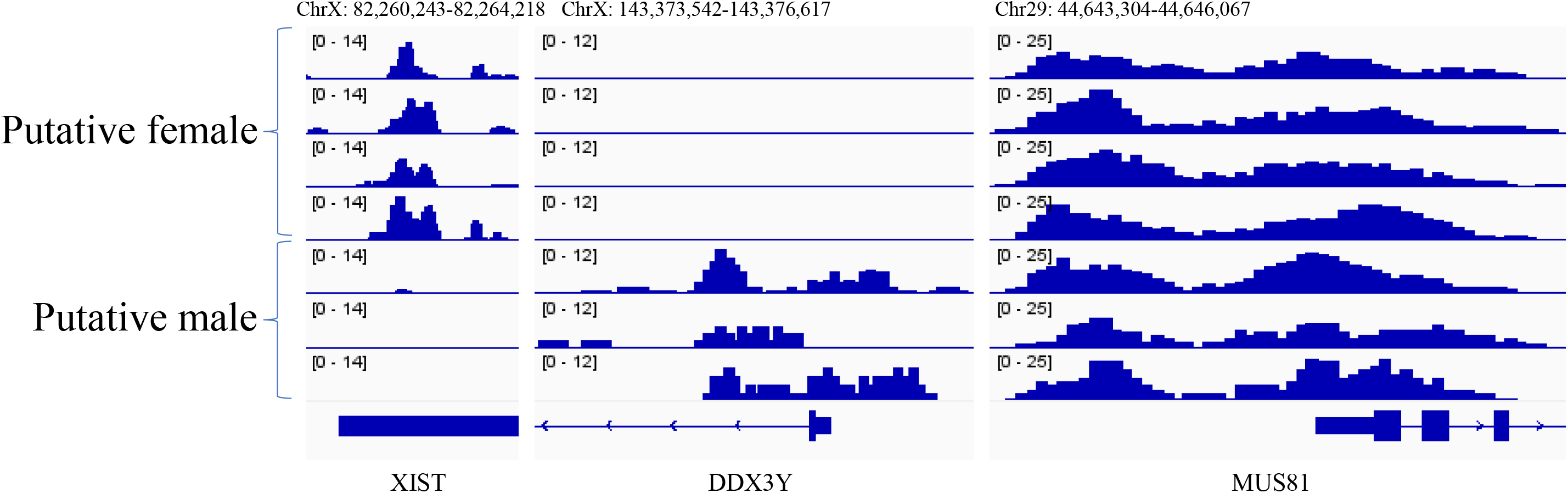
H3K4me3 modifications at *XIST* and *DDX3Y* as male- and female-specific markers, respectively, from NGS-based WOW-CAT. *MUS81* is an internal control that shows stable sex independent H3K4me3 modifications.

We first checked how well the results from the conventional sex identification method (23) matched those from histone modification-based sex identification using WOW-CAT and subsequent qPCR with the 18 embryos remaining after TE cell biopsy (Fig. 6a). The conventional sex identification method (23) determined 7 as male and 11 as female. In qPCR, *MUS81* was selected as an internal control to indicate experimental stability when it showed a positive amplification curve and stable melt curve peak temperature at approximately 81℃. *DDX3Y* was positively determined by the appearance of an amplification curve and stable melt curve with a peak temperature of approximately 81℃, while it was considered undetermined in the absence of an amplification plot or presence of an unstable melt curve. Taking these criteria, 6 out of 7 (86%) male and all 11 (100%) female embryos were identified as such also in WOW-CAT-qPCR for *DDX3Y* (total matching was 17/18 = 94.4%) (Fig. 6b). For *XIST*, setting the threshold for the rise of the amplification curve to 30 cycles with a melt curve peaking at 78℃ resulted in 100% concordance (17/17, one male embryo was used in another experiment and was excluded from the calculation) (Fig. 6c).

**Figure 6.**
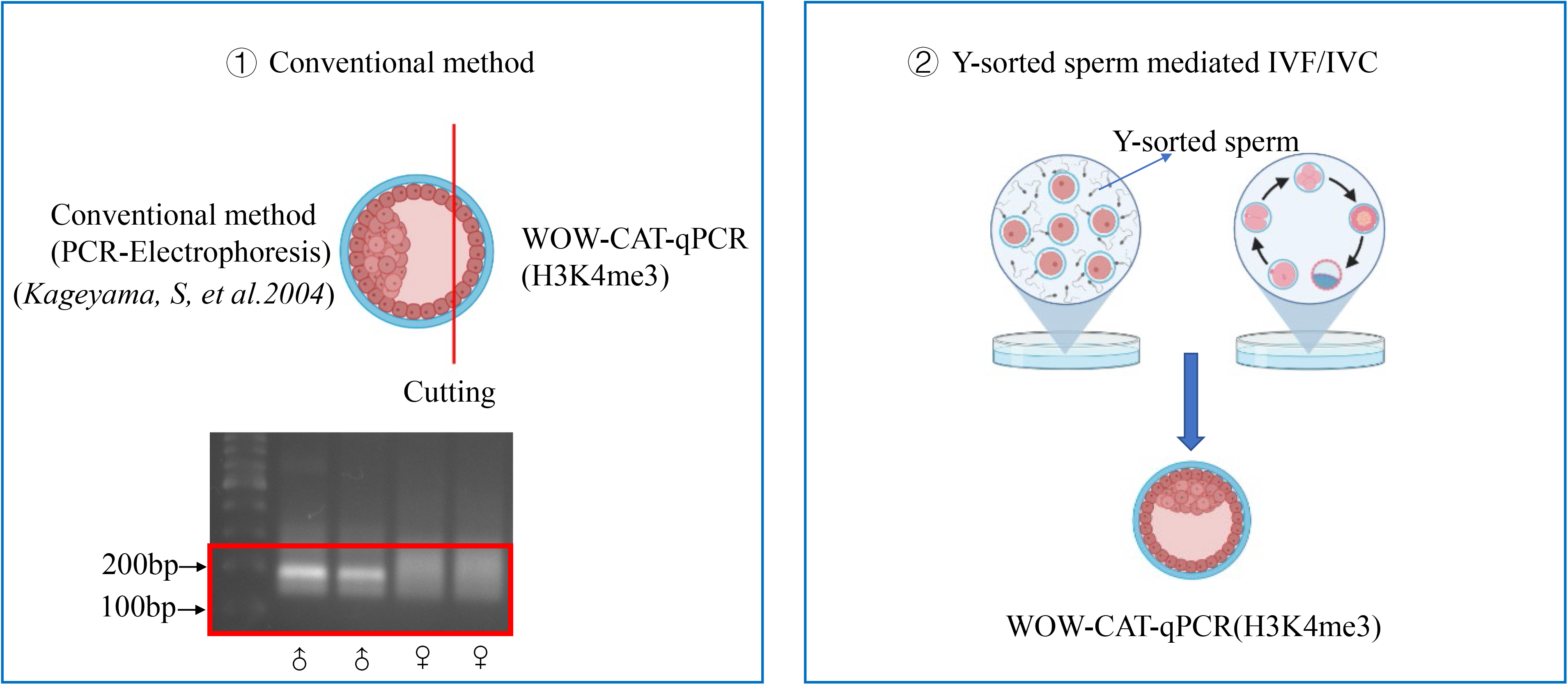
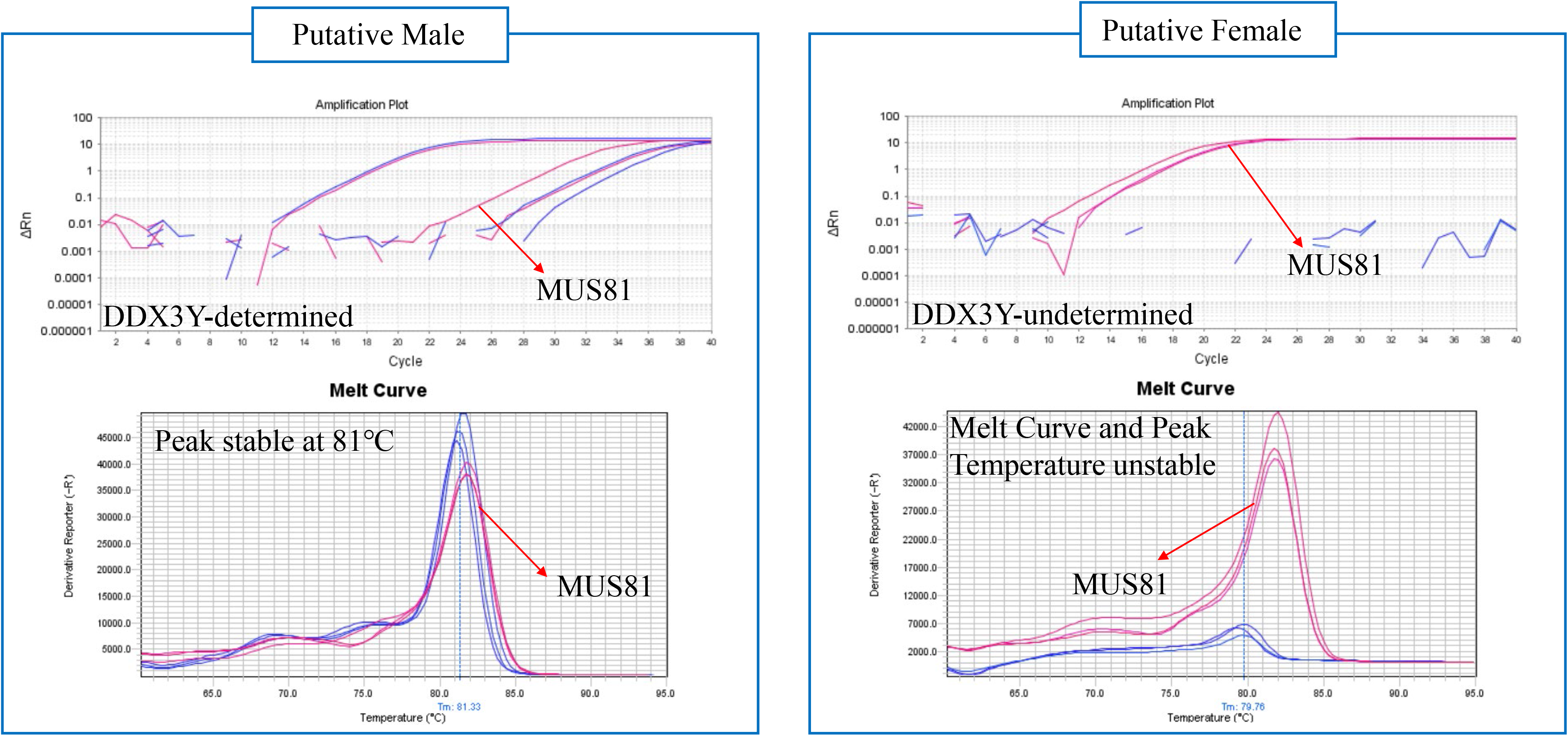
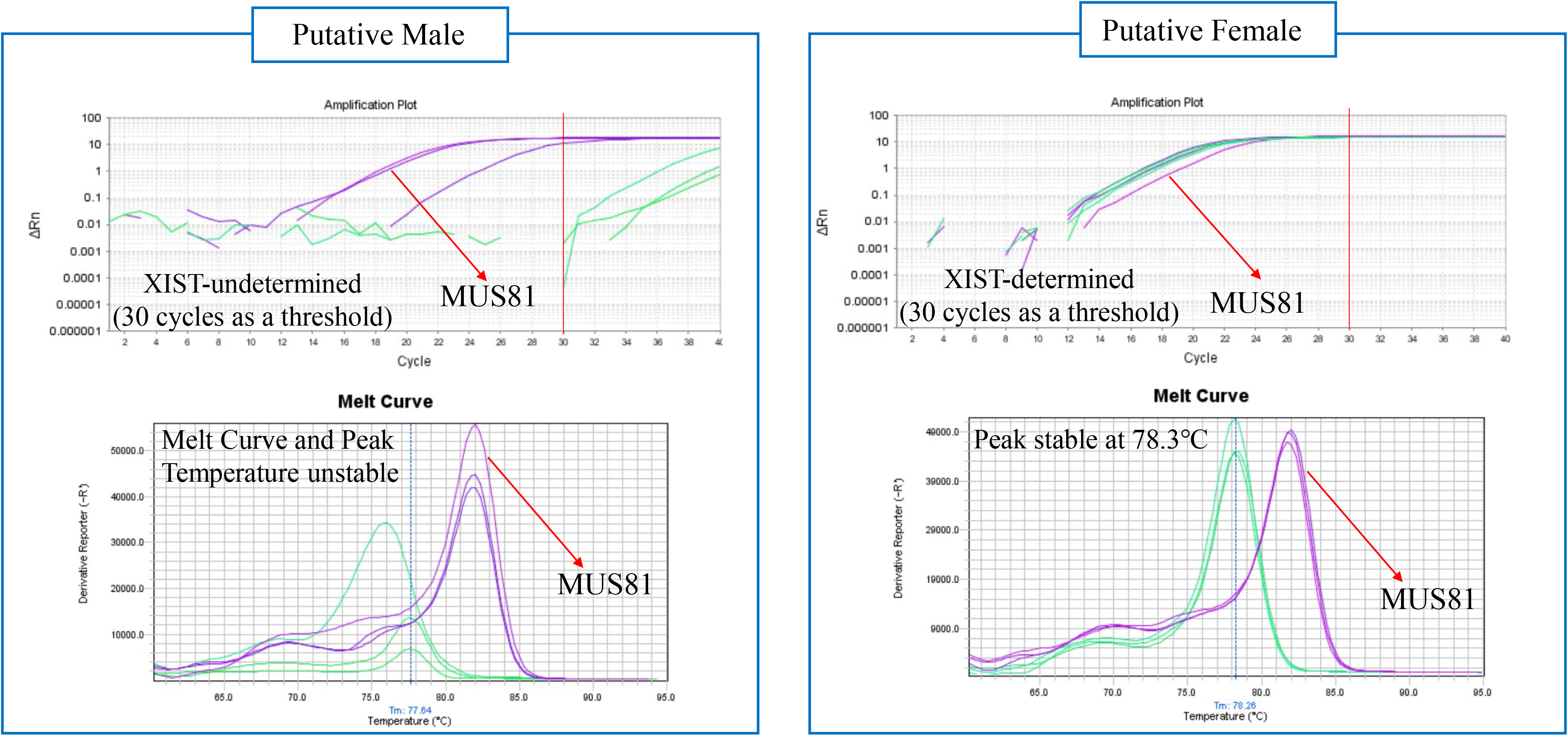
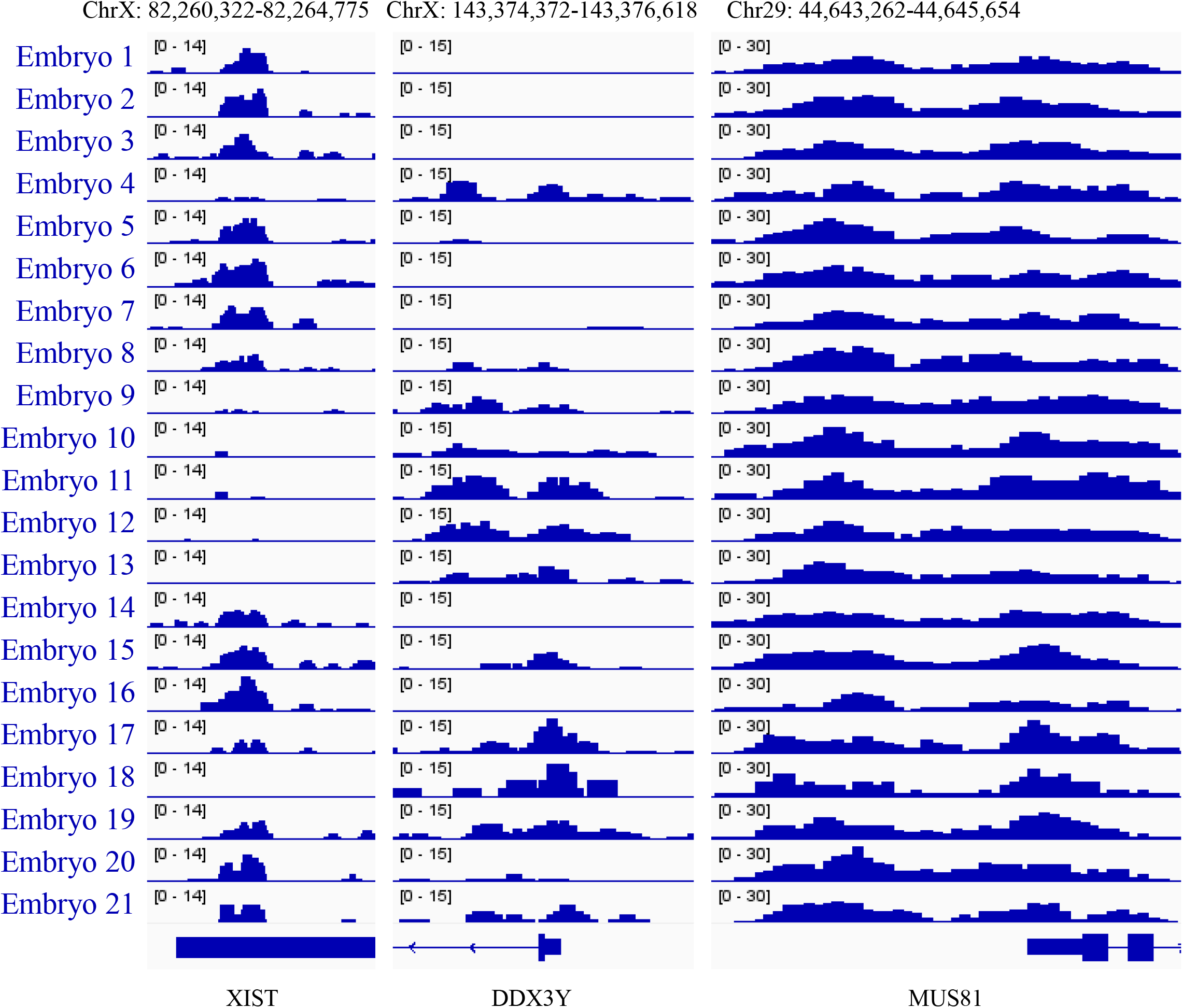
Sex identification accuracy validation. (a) Schema of comparison between ① the conventional sex identification method and ② IVF using Y chromosome-sorted sperm and WOW- CAT-qPCR-based sex identification. All icons except the electrophoresis picture are from Biorender.com. (b) Comparison of the results between the conventional sex identification method and WOW-CAT-qPCR for *DDX3Y*. (c) Comparison between the conventional sex identification method and WOW-CAT-qPCR for *XIST*. For the amplification plot, threshold for the rise of the curve was set to 30 cycles. (d) NTU-CAT results at *XIST* and *DDX3Y* in multiple blastocysts.

In addition, we assessed the embryos derived from IVF using Y chromosome-sorted sperm. WOW-CAT-qPCR identified 10 embryos out of 12 (83.3%) as male using *DDX3Y* as a marker, consistent with the approximately 90% sex ratio of Y chromosome-sorted sperm. In contrast, *XIST* sometimes showed unexpected amplification from presumptive male (*DDX3Y*+) samples, which was consistent with the results of the NGS data (H3K4me3 modifications in both genes) in NTU-CAT (Fig. 6d). Thus, the *XIST* marker did not seem to be reliable for screening whole embryo-derived WOW- CAT DNA libraries. We assessed the biopsied/main parts of the NGS-based WOW-CAT results again and found that the biopsied (TE) parts showed nearly no modification of *XIST* in putative male (*DDX3Y*+) embryos while their main parts unexpectedly harbored *XIST* modifications (Fig. 7). Therefore, the Y chromosome-sorted sperm-derived embryos (using whole embryos) may not be suitable to verify the efficacy of WOW-CAT-qPCR using *XIST* as a marker because some male embryos showed modifications of not only *DDX3Y* but also *XIST*, particularly in the ICM-containing part of the embryo. As assumed, there was no correlation between *XIST* marker-based WOW-CAT- qPCR sex identification and expected sex (male) derived from Y chromosome-sorted sperm (data not shown).

**Figure 7.**
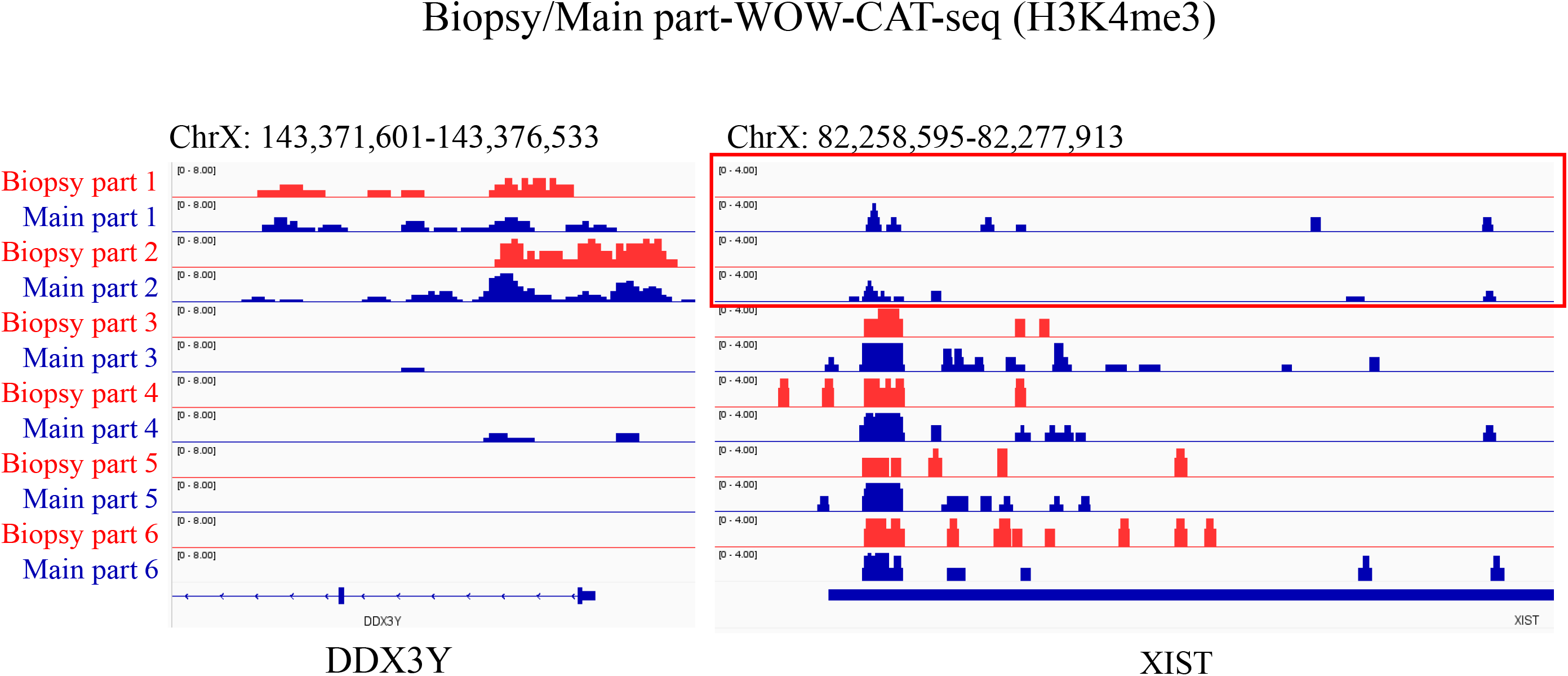
NGS-based WOW-CAT results at *XIST* and *DDX3Y* in the biopsied and main parts of blastocysts.

## Discussion

The conventional CUT&Tag method (14, 15) can analyze histone modifications in a small number of cells. This approach utilizes concanavalin A-coated magnetic beads to fix dispersed cells to the solid phase in order to facilitate handling and processing of the cells. However, preimplantation embryos are cell masses that can be individually transferred to any reaction solution in the experimental process using a fine pipette. Recently, the new method of NTU-CAT (16) was developed that can handle embryos with good permeability of antibodies even without a solid phase. With the successful implementation of NTU-CAT, we further modified it by taking advantage of the WOW system (17, 18), which enables the processing of multiple embryos in a shorter time with less reagent. Because WOW-CAT required only liquid exchange without embryo transfer until the tagmentation step, systematic errors and cell loss were reduced and facilitated the profiling of histone modifications using not only a single blastocyst but also a small part of it.

As a result, WOW-CAT generated genome-wide profiles of representative H3K4me3 histone modifications from single embryos, which were comparable to the results obtained using the conventional ChIP-seq method and NTU-CAT. We demonstrated the overall similarity of the signals detected among these three methods (Fig. 2a–c). The shape of the average profile of the peaks near the TSSs of genes differed among the three methods, such that the “valleys” detected by ChIP-seq were not observed with NTU-CAT but were slightly detected with WOW-CAT (Fig. 2b). The difficulty in detecting “valleys” is a general feature of CUT&Tag experiments (15, 31) and may be due to the bias of Tn5 transposase to preferentially cut open chromatin regions (32). The magnitude of this bias is estimated to generate 10%–15% false positive peaks per sample, which are detected possibly due to the open chromatin bias of Tn5 transposase (16). The valley-like trend of WOW-CAT, which indicates an intermediate form between ChIP-seq and NTU-CAT, suggests that WOW-CAT may alleviate this bias of Tn5 transposase. Furthermore, high correlations of the signals per genomic bin were obtained between WOW-CAT and ChIP-seq (Pearson correlation = 0.88–0.91), which showed nearly no difference between NTU-CAT and ChIP-seq (Pearson correlation = 0.89–0.91), between WOW-CAT and NTU-CAT (Pearson correlation = 0.95–0.97), and within the WOW-CAT replicates (Pearson correlation = 0.96–0.97, Fig. 2c).

The reduction of systematic errors by WOW-CAT compared with NTU-CAT made it possible to obtain results from a cell mass smaller than a whole blastocyst. Then, simultaneous WOW-CAT of two equal parts of the embryo was conducted for H3K4me3 and H3K27ac, respectively. Fig. 3a shows the same region as reported by Zhou et al. (30), who used 1,000–1,500 blastomeres for CUT&Tag to detect these modifications. A high similarity was demonstrated in the common (unshaded), H3K4me3- dominant (blue-shaded), and H3K27ac-dominant (red-shaded) areas, indicating a high similarity between conventional CUT&Tag using a group of embryos and WOW-CAT using even a part of single embryos. The average profile plots of the H3K4me3/H3K27ac signals around the TSSs are shown in Fig. 3b. The highly similar valley-like trend might be due to the same gene expression-promoting function of H3K4me3 and H3K27ac. The H3K4me3 signals around the TSSs were higher than those of H3K27ac, which is also consistent with the results shown by Zhou et al., who used the same antibodies (30). This is the first simultaneous detection of H3K4me3 and H3K27ac modifications within identical single blastocysts. Fig. 3c and d show the heatmap and volcano plot, respectively, of the normalized H3K4me3 and H3K27ac modifications within three single embryos. Among the 2,500 genes analyzed, 148 and 123 genes were H3K4me3- and H3K27ac-dominant, respectively, and the other genes showed a balanced state. Intriguingly, the H3K4me3- and H3K27ac-dominant genes are involved in different biological processes. For example, the protein encoded by the H3K4me3- dominant gene *NDUFA3* is a subunit of ubiquinone, which is located in the mitochondrial inner membrane and is the largest of the five members of the electron transport chain (33). The protein encoded by the H3K27ac-dominant gene *GJB3* is a component of gap junctions, which provide a route for the cell-to-cell diffusion of low molecular weight materials (34). Although H3K4me3 and H3K27ac are both gene expression-promoting modifications, this is the first time that subtle differences in their dominance in different genes and biological processes at the level of an identical blastocyst have been shown (Fig. 3d–f).

We next attempted to biopsy a small part of a single blastocyst for WOW-CAT to establish a method for diagnosing the properties of embryos. We biopsied only TE cells to preserve the integrity of the blastocysts for future embryo transfer. The precision of TE cell biopsy was examined (Fig. 4a) and the high rate of average CDX2/Hoechst-stained cells (21 out of 22, 95%) in the biopsied part indicated that the TE cells were sampled precisely. In addition, the absence of H3K4me3 modifications of *NANOG* (an ICM-specific marker) in the biopsied part further showed the precision of TE cell biopsy (Fig. 4b). A snapshot of the WOW-CAT H3K4me3 peaks of the whole, biopsied, and remaining (main) parts of blastocysts is shown in Fig. 4c. The overall landscape depicted by the location and shape of the peaks was almost the same among them. This indicated that the H3K4me3 modification level of TE cells can generally reflect that of the remaining part of the blastocyst, suggesting the possible usage of TE (biopsy) cell modifications for diagnosis. With the successful construction of DNA libraries for the enrichment of specific histone modifications from biopsied TE cells by WOW-CAT, we developed a qPCR-based detection protocol for important histone markers from TE cell-derived DNA libraries, which directly reflected the properties of the remaining part of the blastocyst.

We selected H3K4me3 modifications at *XIST* and *DDX3Y* as possible sex-identification markers for females and males, respectively, from the NGS-based WOW-CAT analysis (Fig. 5), in which embryos generally showed only one modification, either *XIST* (activated only in females) or *DDX3Y* (a Y chromosome gene), suggesting they were female and male embryos, respectively.

Then, we validated the accuracy of the WOW-CAT-qPCR method for sex identification by comparing it with the conventional sex identification method (23) and by using Y chromosome-sorted sperm-derived embryos. The results for *XIST* and *DDX3Y* were well matched with those of the conventional sex identification method, but only *DDX3Y* matched well in the Y chromosome-sorted sperm-based method (Fig. 6a–c). This may be explained by the fact that we only used the biopsied (TE) part in the comparison with the conventional sex identification method, which was applied to the remaining main part of the same embryos in the present study, while we used whole blastocysts in the comparison with the Y chromosome-sorted sperm-based experiment. Thus, we checked the NGS results of WOW-CAT for the biopsied and main parts again and found that the main part containing the ICM had a faint *XIST* modification, despite the presence of the *DDX3Y* modification, while the TE part was completely free of the *XIST* modification within the same embryos. (Fig. 6d). Therefore, the comparison with the conventional sex identification method was considered to verify the accuracy of WOW-CAT-qPCR, but the Y chromosome-sorted sperm-derived embryos may not be suitable for this purpose. In addition, this diagnostic method was designed to use the TE part to preserve the integrity of blastocysts for future embryo transfer. Thus, it does not matter that *XIST* sometimes shows unexpected amplification in presumptive male (*DDX3Y*+) samples (Fig. 5a, b). However, the reason for the inconsistency of the modification level between the TE part and remaining part in male blastocysts is unclear and more research is required to explain this interesting phenomenon.

As we successfully biopsied the TE cells for WOW-CAT and obtained *XIST/DDX3Y* amplification from qPCR, we consider that a basic diagnostic method for embryos was established, at least in terms of sex identification. It is anticipated that the identification of useful epigenetic modifications will continue to progress for histone modifications, which will allow for the evaluation of the quality of embryos beyond sex (6). If useful markers are identified, they may be used for the quality control of embryos themselves and embryo production protocols contributing to improved embryo quality in assisted reproductive technologies (ART).

We used a bovine model because bovine embryos are more similar to human embryos than rodent models in many respects, including mono-ovulatory nature, gamete size, embryonic developmental speed, blastocyst cell numbers, and the timing of embryonic genome activation, which makes it a clinically important model for the study of human embryos (35). However, further validation in other experimental animal models is needed to explore a wider range of applications for WOW-CAT.

## Conclusion

WOW-CAT enables the profiling of genome-wide histone modifications from not just a single blastocyst but also from a portion of it. By using this method, histone modifications were analyzed in two halves of the same single blastocyst and in the TE cell part for the first time. By combining WOW- CAT with qPCR for the TE cell part, information on specific histone markers can be detected, which reflect the properties of the remaining part of the embryo without the need for deep sequencing. These results suggest the applicability of WOW-CAT for flexible epigenetic analysis in individual embryos as well as for preimplantation epigenetic diagnosis. With the anticipation of discovering markers indicative of embryo quality, this procedure can be further perfected to ensure embryo quality control before transfer, which will be significant for improving the efficacy of ART.

## Notes

### Competing Interest Statement

The authors have declared no competing interest.

## References

1. Zentner GE, Henikoff S. High-resolution digital profiling of the epigenome. Nat Rev Genet. 2014;15(12):814–27.

2. Li G, Yu Y, Fan Y, Li C, Xu X, Duan J, et al. Genome wide abnormal DNA methylome of human blastocyst in assisted reproductive technology. J Genet Genomics. 2017;44(10):475–81.

3. Olcha M, Dong X, Feil H, Hao X, Lee M, Jindal S, et al. A workflow for simultaneous DNA copy number and methylome analysis of inner cell mass and trophectoderm cells from human blastocysts. Fertil Steril. 2021;115(6):1533–40.

4. Yang M, Tao X, Scott K, Zhan Y, Scott RT, Seli E. Evaluation of genome-wide DNA methylation profile of human embryos with different developmental competences. Hum Reprod. 2021;36(6):1682–90.

5. Yu B, Smith TH, Battle SL, Ferrell S, Hawkins RD. Superovulation alters global DNA methylation in early mouse embryo development. Epigenetics. 2019;14(8):780–90.

6. Bai D, Sun J, Chen C, Jia Y, Li Y, Liu K, et al. Aberrant H3K4me3 modification of epiblast genes of extraembryonic tissue causes placental defects and implantation failure in mouse IVF embryos. Cell Rep. 2022;39(5):110784.

7. Lu X, Zhang Y, Wang L, Wang L, Wang H, Xu Q, et al. Evolutionary epigenomic analyses in mammalian early embryos reveal species-specific innovations and conserved principles of imprinting. Sci Adv. 2021;7(48):eabi6178.

8. Dahl JA, Jung I, Aanes H, Greggains GD, Manaf A, Lerdrup M, et al. Broad histone H3K4me3 domains in mouse oocytes modulate maternal-to-zygotic transition. Nature. 2016;537(7621):548–52.

9. Liu X, Wang C, Liu W, Li J, Li C, Kou X, et al. Distinct features of H3K4me3 and H3K27me3 chromatin domains in pre-implantation embryos. Nature. 2016;537(7621):558–62.

10. Zhang B, Zheng H, Huang B, Li W, Xiang Y, Peng X, et al. Allelic reprogramming of the histone modification H3K4me3 in early mammalian development. Nature. 2016;537(7621):553–7.

11. Schmid M, Durussel T, Laemmli UK. ChIC and ChEC; genomic mapping of chromatin proteins. Mol Cell. 2004;16(1):147–57.

12. Skene PJ, Henikoff S. An efficient targeted nuclease strategy for high-resolution mapping of DNA binding sites. Elife. 2017;6.

13. Skene PJ, Henikoff JG, Henikoff S. Targeted in situ genome-wide profiling with high efficiency for low cell numbers. Nat Protoc. 2018;13(5):1006–19.

14. Kaya-Okur HS, Janssens DH, Henikoff JG, Ahmad K, Henikoff S. Efficient low-cost chromatin profiling with CUT&Tag. Nat Protoc. 2020;15(10):3264–83.

15. Kaya-Okur HS, Wu SJ, Codomo CA, Pledger ES, Bryson TD, Henikoff JG, et al. CUT&Tag for efficient epigenomic profiling of small samples and single cells. Nat Commun. 2019;10(1):1930.

16. Susami K, Ikeda S, Hoshino Y, Honda S, Minami N. Genome-wide profiling of histone H3K4me3 and H3K27me3 modifications in individual blastocysts by CUT&Tag without a solid support (NON-TiE-UP CUT&Tag). Sci Rep. 2022;12(1):11727.

17. Vajta G, Peura TT, Holm P, Paldi A, Greve T, Trounson AO, et al. New method for culture of zona-included or zona-free embryos: the Well of the Well (WOW) system. Mol Reprod Dev. 2000;55(3):256–64.

18. Vajta G, Korosi T, Du Y, Nakata K, Ieda S, Kuwayama M, et al. The Well-of-the-Well system: an efficient approach to improve embryo development. Reprod Biomed Online. 2008;17(1):73–81.

19. Ishibashi M, Ikeda S, Minami N. Comparative analysis of histone H3K4me3 modifications between blastocysts and somatic tissues in cattle. Sci Rep. 2021;11(1):8253.

20. Yamazaki S, Ikeda S, Minami N. Comparative analysis of histone H3K27me3 modifications between blastocysts and somatic tissues in cattle. Anim Sci J. 2022;93(1):e13684.

21. de Sousa RV, da Silva Cardoso CR, Butzke G, Dode MAN, Rumpf R, Franco MM. Biopsy of bovine embryos produced in vivo and in vitro does not affect pregnancy rates. Theriogenology. 2017;90:25–31.

22. Wydooghe E, Vandaele L, Beek J, Favoreel H, Heindryckx B, De Sutter P, et al. Differential apoptotic staining of mammalian blastocysts based on double immunofluorescent CDX2 and active caspase-3 staining. Anal Biochem. 2011;416(2):228–30.

23. Kageyama S, Yoshida I, Kawakura K, Chikuni K. A novel repeated sequence located on the bovine Y chromosome: its application to rapid and precise embryo sexing by PCR. J Vet Med Sci. 2004;66(5):509–14.

24. Hirayama H, Kageyama S, Moriyasu S, Sawai K, Onoe S, Takahashi Y, et al. Rapid sexing of bovine preimplantation embryos using loop-mediated isothermal amplification. Theriogenology. 2004;62(5):887–96.

25. Langmead B, Salzberg SL. Fast gapped-read alignment with Bowtie 2. Nat Methods. 2012;9(4):357–9.

26. Shen L, Shao N, Liu X, Nestler E. ngs.plot: Quick mining and visualization of next-generation sequencing data by integrating genomic databases. BMC Genomics. 2014;15:284.

27. Robinson JT, Thorvaldsdottir H, Winckler W, Guttman M, Lander ES, Getz G, et al. Integrative genomics viewer. Nat Biotechnol. 2011;29(1):24–6.

28. Kaya-Okur HS, Wu SJ, Codomo CA, Pledger ES, Bryson TD, Henikoff JG, et al. CUT&Tag for efficient epigenomic profiling of small samples and single cells. Nature Communications. 2019;10(1):1930.

29. Fujiwara Y, Tanno Y, Sugishita H, Kishi Y, Makino Y, Okada Y. Preparation of optimized concanavalin A-conjugated Dynabeads(R) magnetic beads for CUT&Tag. PLoS One. 2021;16(11):e0259846.

30. Zhou C, Halstead MM, Bonnet-Garnier A, Schultz RM, Ross PJ. Resetting H3K4me3, H3K27ac, H3K9me3 and H3K27me3 during the maternal-to-zygotic transition and blastocyst lineage specification in bovine embryos. 2022:2022.04.07.486777.

31. Fujiwara Y, Tanno Y, Sugishita H, Kishi Y, Makino Y, Okada Y. Preparation of optimized concanavalin A-conjugated Dynabeads® magnetic beads for CUT&Tag. PLoS One. 2021;16(11):e0259846.

32. Wang M, Zhang Y. Tn5 transposase-based epigenomic profiling methods are prone to open chromatin bias. 2021:2021.07.09.451758.

33. Rak M, Rustin P. Supernumerary subunits NDUFA3, NDUFA5 and NDUFA12 are required for the formation of the extramembrane arm of human mitochondrial complex I. FEBS letters. 2014;588(9):1832–8.

34. Huo Y, Zhou Y, Zheng J, Jin G, Tao L, Yao H, et al. GJB3 promotes pancreatic cancer liver metastasis by enhancing the polarization and survival of neutrophil. Frontiers in immunology. 2022;13:983116.

35. Ménézo YJ, Hérubel F. Mouse and bovine models for human IVF. Reprod Biomed Online. 2002;4(2):170–5.

